# A neutrophil - B-cell axis governs disease tolerance during sepsis via Cxcr4

**DOI:** 10.1101/2022.03.21.485114

**Authors:** Riem Gawish, Barbara B. Maier, Georg Obermayer, Martin L. Watzenböck, Anna-Dorothea Gorki, Federica Quattrone, Asma Farhat, Karin Lakovits, Anastasiya Hladik, Ana Korosec, Arman Alimohammadi, Ildiko Mesteri, Felicitas Oberndorfer, Fiona Oakley, John Brain, Louis Boon, Irene Lang, Christoph J. Binder, Sylvia Knapp

## Abstract

Sepsis is a life-threatening condition characterized by uncontrolled systemic inflammation and coagulation, leading to multi-organ failure. Therapeutic options to prevent sepsis-associated immunopathology remain scarce.

Here, we established a model of long-lasting disease tolerance during severe sepsis, manifested by diminished immunothrombosis and organ damage in spite of a high pathogen burden. We found that, both neutrophils and B cells emerged as key regulators of tissue integrity. Enduring changes in the transcriptional profile of neutrophils, included upregulated Cxcr4 expression in protected, tolerant hosts. Neutrophil Cxcr4 upregulation required the presence of B cells, suggesting that B cells promoted tissue tolerance by suppressing tissue damaging properties of neutrophils. Finally, therapeutic administration of a Cxcr4 agonist successfully promoted tissue tolerance and prevented liver damage during sepsis. Our findings highlight the importance of a critical B-cell/neutrophil interaction during sepsis and establish neutrophil Cxcr4 activation as a potential means to promote disease tolerance during sepsis.

**Summary:** We show that a B cell/neutrophil interaction in the bone marrow facilitates tissue tolerance during severe sepsis. By affecting neutrophil Cxcr4 expression, B cells can impact neutrophil effector functions. Finally, therapeutic activation of Cxcr4 successfully promoted tissue tolerance and prevented liver damage during sepsis.

## Introduction

Sepsis is a life-threatening condition triggered by severe infections with bacteria, viruses or fungi. In spite of the successful use of antimicrobial therapies, mortality rates remain high with up to 50%, (1, 2). The main determinant of sepsis-associated mortality is rarely the pathogen, but instead the combination of dysregulated systemic inflammation, immune paralysis and hemostatic abnormalities that together cause multi-organ failure (3). Upon pathogen sensing, ensuing inflammation promotes the activation of coagulation, which in turn generates factors that further amplify inflammation, thus creating a vicious, self-amplifying cycle. These events result in systemic inflammation and the widespread formation of microvascular thrombi that together cause vascular leak, occlusion of small vessels and ultimately multi-organ failure (4, 5). Whether a patient suffering from sepsis enters this fatal circuit of immunopathology or instead is able to maintain vital organ functions and survives sepsis is not well understood (6–8).

The concept of “disease tolerance” describes a poorly studied, yet essential host defense strategy, on top of the well understood strategies of avoidance and resistance (9). While avoidance means preventing pathogen exposure and infection, and resistance aims to more efficiently reduce the pathogen load in the course of an established infection, disease tolerance involves mechanisms, which minimize the detrimental impact of infection-associated immunopathology, thus improving host fitness despite the infection (9, 10). To this end, a number of mechanisms that shape the process of disease tolerance have been suggested, including alterations in cellular metabolism, DNA damage response, tissue remodeling or oxidative stress (10). However, little is known about the specific contribution of immune cells to disease tolerance during severe infections, and therapeutic options to increase disease tolerance are limited due to a lack of knowledge about detailed molecular and cellular tolerance mechanisms (6–8).

In this study, we investigated mechanisms of disease tolerance, by comparing tolerant and sensitive hosts during a severe bacterial infection. While sensitive animals developed severe coagulopathy and tissue damage during sepsis, tolerant animals were able to maintain tissue integrity in spite of a high bacterial load. Tolerance was induced by the prior exposure of animals to a single, low-dose of LPS and could be uncoupled from LPS-induced suppression of cytokine responses. We provide evidence for a deleterious and organ-damaging interaction between B cells and neutrophils during sepsis in sensitive animals, while in tolerant animals neutrophils and B cells jointly orchestrated tissue protection during sepsis, which was associated with transcriptional reprogramming of neutrophils and B cell dependent upregulation of neutrophil Cxcr4. Our data suggest that B cells can modulate the tissue damaging properties of neutrophils by influencing neutrophil Cxcr4 signaling. Consequently, the administration of a Cxcr4 agonist prevented sepsis-associated microthrombosis and resulting tissue damage, thereby exposing a potential therapeutic strategy to foster tissue tolerance in severe sepsis.

## Results

### LPS pre-exposure induces long-term tissue tolerance during Gram-negative sepsis

To establish a model for the study of tissue tolerance during sepsis we challenged mice intravenously (i.v.) with a subclinical dose of LPS 1 day, 2 weeks, 5 weeks or 8 weeks, respectively, prior to the induction of Gram-negative sepsis by intraperitoneal (i.p.) injection of the virulent *E. coli* strain O18:K1. While LPS pretreatment 24h prior to infection significantly improved pathogen clearance, any longer period (i.e. 2-8 weeks) between LPS administration and infection did not affect the bacterial load when compared to control mice (Figure 1A, Figure S1A). Importantly though, all LPS pre-treated groups were substantially protected from sepsis-associated tissue damage, illustrated by the absence of elevated liver transaminase (ASAT and ALAT) plasma levels (Figure 1B). Thus, short-term (24h) LPS pre-exposure improved resistance to infection and consequently tissue integrity, while long-term (2-8 weeks) LPS pre-exposure enabled the maintenance of tissue integrity irrespective of a high bacterial load, which per definition resembles disease tolerance.

**Figure 1:**
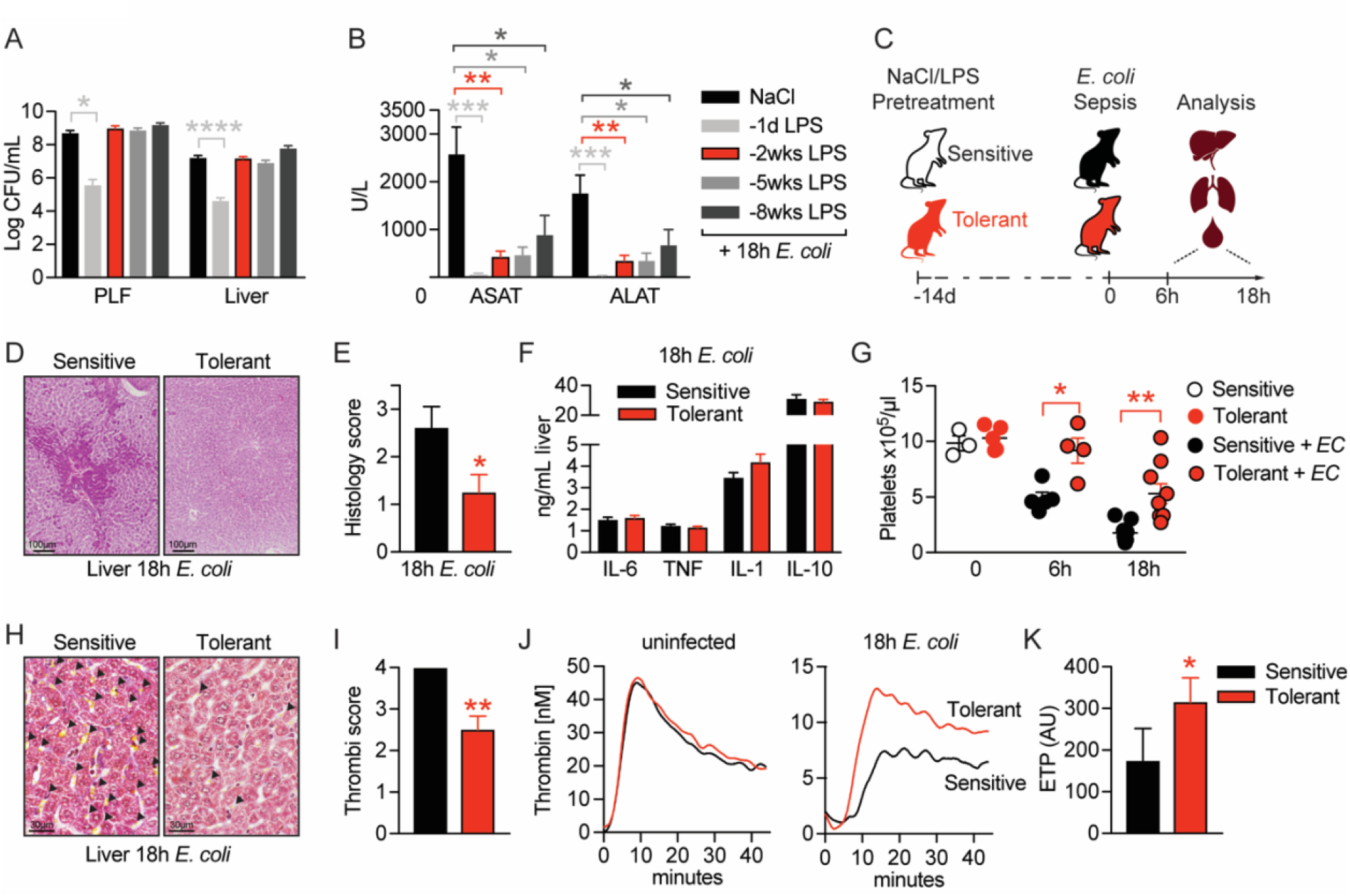
LPS pre-exposure induces long-term tissue tolerance during Gram-negative sepsiss. (**A**) *E. coli* CFU 18h p.i. in peritoneal lavage fluid (PLF) and liver of mice, which were pretreated with LPS or NaCl at depicted time points before infection. (**B**) ASAT and ALAT plasma levels at 18h p.i. of mice, pretreated with LPS or NaCl at indicated time points. (**C**) Schematic depiction of the treatment procedure and endpoints. (**D-E**) H&E staining and pathology scores of liver sections from mice pretreated with LPS or NaCl 2 weeks earlier, and infected with *E. coli* for 18h. (**F**) Liver cytokine levels at 18h p.i. (**G**) Blood platelet counts at 2 weeks post LPS/NaCl pretreatment, and 6h or 18h post infection. (**H-I**) Martius, Scarlet and Blue (MSB) fibrin staining of liver sections and scoring of liver microthrombi at 18h p.i. (**J**) *In vitro* thrombin generation capacity of plasma from LPS/NaCL pretreated uninfected and infected mice. (**K**) Endogenous thrombin potential (ETP) of plasma samples 18h p.i. Data in (A) and (B) are pooled from 2-3 independent experiments (n = 6-7/experimental group). Data in (G) are pooled from 2 independent experiments (n = 1-3/experimental group for uninfected and 4-5 for infected mice). All other data are representative for two or more independent experiments (n = 8/experimental group). All data are and presented as mean +/− SEM. * p ≤ 0.05, ** p ≤ 0.01, *** p ≤ 0.001 and **** p ≤ 0.0001.

To dissect the underlying mechanism of tissue tolerance, we thus performed all subsequent experiments by treating mice with either LPS or saline two weeks prior to bacterial infection, allowing us to compare tolerant with sensitive hosts. Mice were either sacrificed two weeks after LPS pretreatment to assess changes in tolerant hosts prior to infection, or six to 18h after *E. coli* infection to determine early (6h) or late inflammation and organ damage (18h), respectively, during sepsis (Fig. 1C). Doing so, we observed that organ protection (Figure 1B) was associated with the absence of liver necrosis (Figure 1D and 1E), while inflammatory cytokine and chemokine levels were indistinguishable between sensitive and tolerant mice 18h post infection (p.i.) (Figure 1F, Figure S1B and 1C). A major cause of organ damage during sepsis is the disseminated activation of coagulation, which is characterized by systemic deposition of micro-thrombi and substantial platelet consumption, resulting in a critical reduction in tissue perfusion (4–6). While we discovered a severe decline in platelet numbers upon *E. coli* infection in sensitive mice, tolerant mice maintained significantly higher blood platelet counts (Figure 1G) and, in sharp contrast to sensitive animals, showed almost no deposition of micro-thrombi in liver (Figure 1H and 1I) and lung sections (Figure S1D), indicating that tissue tolerance occurred systemic and not organ specific. Considering that LPS exposure itself can impact coagulation factor levels and blood platelet numbers (11, 12), we importantly found similar platelet counts in sensitive and tolerant mice at the onset of *E. coli* infection (2 weeks post LPS) (Figure 1G). In addition, we did not detect any indication for an altered coagulation potential in tolerant mice before sepsis induction, as both groups showed a similar plasma thrombin generation potential prior to infection (Figure 1J left panel, Figure S1E). However, compared to sensitive animals, the thrombin generation capacity was only preserved in tolerant mice after infection (18h p.i.), suggesting that tolerance mechanisms prevented sepsis-associated consumption coagulopathy (Figure 1J right panel and 1K). Taken together, low-dose LPS pretreatment prevented the formation of micro-thrombi and induced a long-lasting state of tissue tolerance during subsequent sepsis.

### B cells regulate tissue tolerance during sepsis independent of early inflammatory responses

Considering the long-term protective effect of LPS pre-exposure in tolerant animals, we next tested the possibility that long-lived immune cells like lymphocytes might regulate tissue tolerance during sepsis. Strikingly, the absence of lymphocytes, as in Rag2 deficient (Rag2^−/−^) animals, already resulted in profoundly reduced liver damage upon bacterial infection of naïve, sensitive mice and fully abrogated further LPS-induced tissue tolerance without affecting the bacterial load (Figure 2A, S2A and S2B). This indicated that lymphocytes on the one hand importantly contributed to the sensitivity of animals to sepsis-associated organ damage in naïve mice, and on the other hand were essential in mediating LPS-induced tissue protection in tolerant hosts. In support of this, adoptive transfer of splenocytes into Rag2^−/−^ mice reestablished LPS-induced tissue tolerance (Figure 2A). Depleting CD8 or CD4 T cells, respectively, prior to LPS exposure (Figure 2B and S2C) neither affected the difference between sensitive and tolerant animals to organ damage (Figure 2C) nor the bacterial load (Figure S2D) upon *E. coli* infection. In contrast, B cell deficiency (J_H_T^−/−^ mice) fully prevented the development of tissue damage during sepsis irrespective of tolerance induction (Figure 2D) and despite a high bacterial load (Figure S2E). Furthermore, adoptive transfer of B cells into Rag2^−/−^ mice (Figure 2E) aggravated liver damage upon *E. coli* infection in sensitive and reestablished tissue protection in tolerant mice (Figure 2F). These findings indicated that B cells, but not T cells, played an ambiguous role as they were involved in both, sepsis-associated organ damage and the establishment of LPS-triggered tissue tolerance. We then tested if splenectomy would replicate the protective effects of B cell deficiency during sepsis and interestingly found that splenectomy was associated with reduced liver damage in naïve, sensitive mice, but, in contrast to complete lymphocyte deficiency, not sufficient to abrogate LPS-induced tissue protection in tolerant animals (Figure 2G and S2F). This suggested that mature splenic B cells contributed to tissue damage during severe infections, while other, not spleen derived, B cell compartments were instrumental in driving disease tolerance.

**Figure 2:**
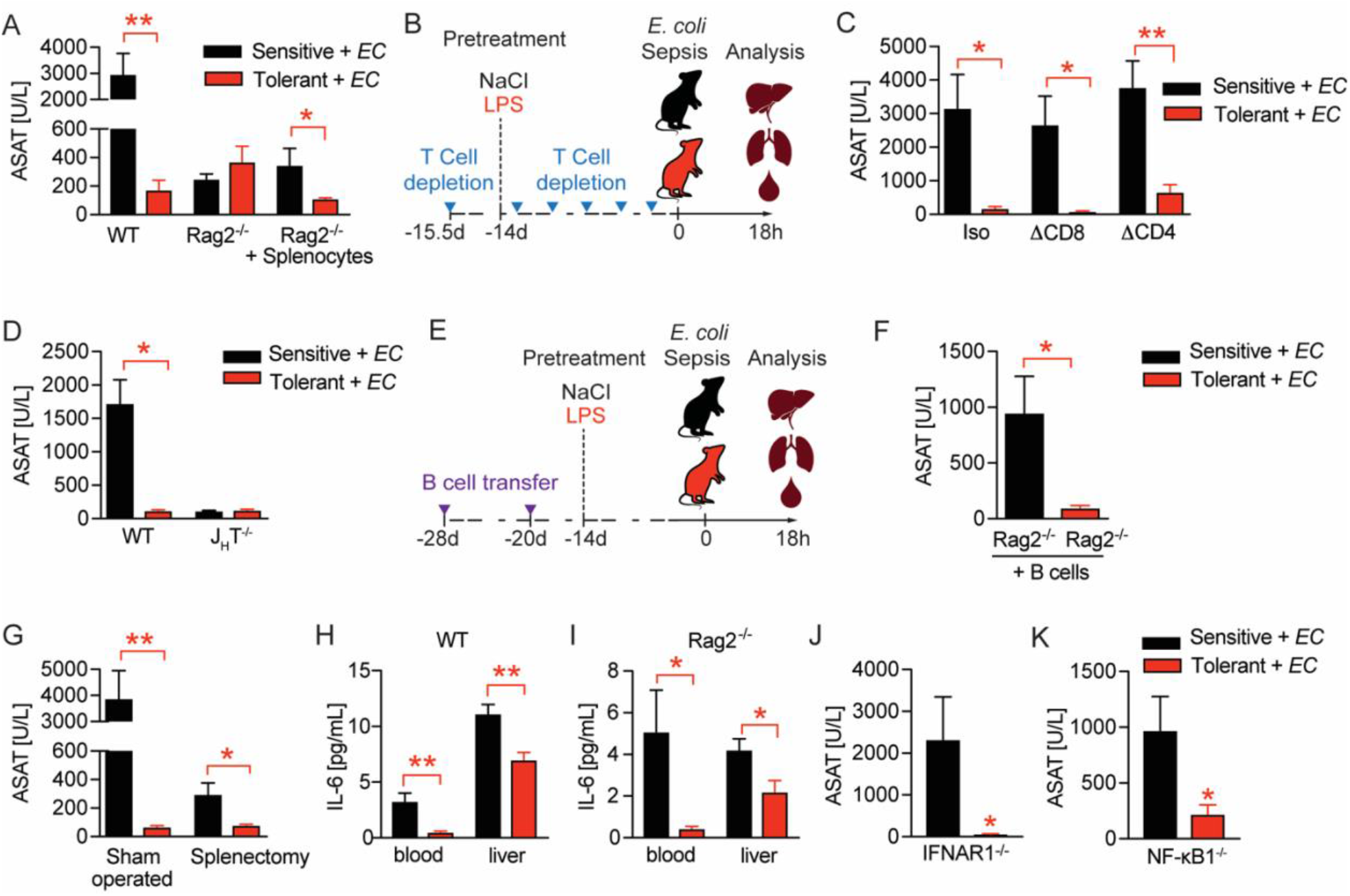
B cells regulate tissue tolerance during sepsis independent of early inflammatory responses. (**A**) ASAT plasma levels at 18h p.i. with *E. coli*, of LPS or NaCl pretreated wildtype or lymphocyte deficient mice (RAG2^−/−^), which have received either PBS or splenocytes i.v. 3 weeks prior infection. (**B**) Schematic depiction of the treatment procedure for T cell depletion experiments. (**C**) ASAT plasma levels 18h p.i. with *E. coli* of mice, which were depleted from CD4^+^ or CD8^+^ T cells prior to LPS or NaCl pretreatment. (**D**) ASAT plasma levels of LPS or NaCl pretreated wildtype or B cell deficient (J_H_T^−/−^) mice at 18h p.i. with *E. coli*. (**E**) Schematic depiction of the treatment procedure for splenocyte and B cell transfer experiments. (**F**) ASAT plasma levels of LPS or NaCl pretreated Rag2^−/−^ mice at 18h p.i. with *E. coli*, which have been reconstituted with bone marrow derived B cells 3 weeks before infection. (**G**) ASAT plasma levels at 18h p.i. with *E. coli* of LPS or NaCl pretreated mice, which were splenectomized or sham operated 1 week before LPS or NaCl pre-exposure (i.e. 3 weeks before infection). (**H**-**I**) IL-6 levels in plasma and liver of NaCl or LPS pretreated wildtype or Rag2^−/−^ mice at 6h p.i. with *E. coli*. (**J**) ASAT plasma levels of NaCl or LPS pretreated IFNAR1^−/−^ mice at 18h p.i. with *E. coli*. (**K**) ASAT plasma levels of NaCl or LPS pretreated NF-κB1^−/−^ mice at 18h p.i. with *E. coli*. Data in (A) and (G-J) and are representative out of 2-3 experiments (n= 3-8/experimental group). Data in (D) and (K) are pooled from 2 independent experiments (n = 2-7/experimental group). Data in (C) and (F) are from a single experiment (n = 6-8/group). Data are and presented as mean +/− SEM. * p ≤ 0.05 and ** p ≤ 0.01.

Given that B cells were shown to promote early production of proinflammatory cytokines such as IL-6 during sepsis in a type I IFN dependent manner (13), we next investigated if LPS pretreatment induced tissue tolerance by dampening B cell driven inflammatory responses. Six hours post *E. coli* infection, we found tolerant wild type mice to exhibit lower IL-6 levels in blood and liver (Figure 2H), as well as lower amounts of important regulators of peritoneal leukocyte migration (14, 15), like CXCL1 and CCL2 (Figure S2G) when compared to sensitive control mice. However, lymphocyte deficient Rag2^−/−^ animals, in whom tissue tolerance could not be induced (Figure 2A and S2A), showed comparable reductions in these mediators of early inflammation in response to LPS pretreatment (Figure 2I and S2H). Further, tolerance during sepsis was induced independent of interferon-α/β receptor (IFNAR) signaling (Figure 2J and S2I) and the anti-inflammatory NF-κB subunit p50 (NF-κB1) (Figure 2K and S2J), which has been shown to mediate the suppression of cytokine production during endotoxin tolerance *in vitro* (16, 17). These data suggested that in tolerant hosts, B cells contributed to tissue protection during sepsis, and that an LPS mediated modulation of early inflammation is unlikely to explain these protective effects.

### Disease tolerance is associated with rearranged B cell compartments

We next compared the B cell compartment in sensitive and tolerant mice and analyzed different B cell populations in spleen and bone marrow 2 weeks after saline or LPS treatment. Tolerance was associated with a mild increase in spleen weight (Figure S3A) and total spleen cell counts, together with an expansion of B cell numbers (Figure 3A). Of note, tolerance-induction by LPS treatment even caused an expansion of transplanted B cells in spleens of Rag2^−/−^ animals (Figure 3B). In parallel, while the total number of bone marrow cells remained indistinguishable (Figure S3B), we found bone marrow B cell numbers increased in tolerant mice, as compared to sensitive controls (Figure 3B). Further analysis of B cell subsets revealed an increase of B1 B cell numbers (IgM^hi^ CD23^−^ CD43^hi^ CD21^−^), a subset that also includes the B1-like innate response activator (IRA) B cells, in spleen and bone marrow (Figure 3D–F and S3C), as well as elevated numbers of pre B and immature and transitional B cells (iB/tB) in the bone marrow (Figure 3F and Figure S3C). In sharp contrast, total numbers of follicular (FO) and marginal zone (MZ) B cells did not change upon tolerance induction (Figure 3D).

**Figure 3:**
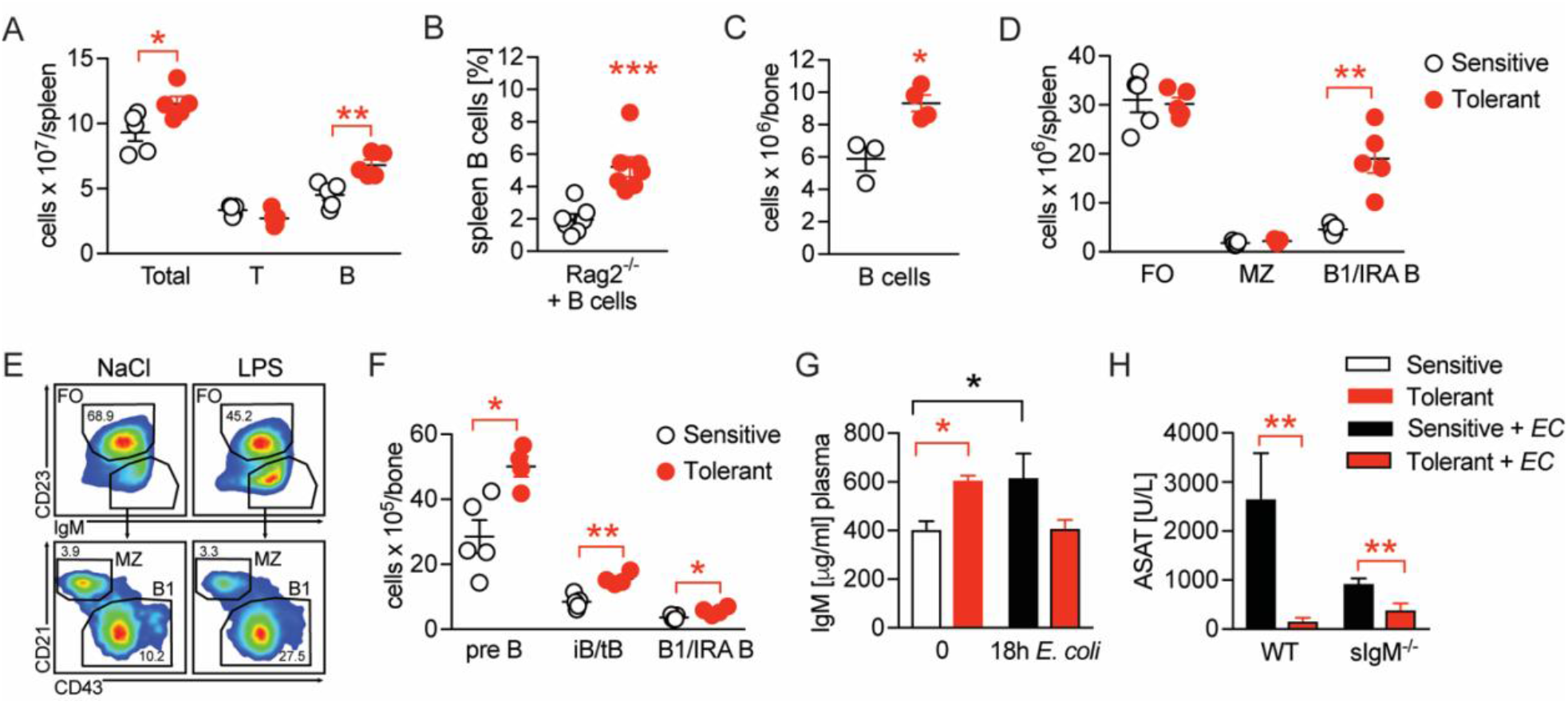
Disease tolerance is associated with rearranged B cell compartments. (**A**) Flow-cytometric analysis of B and T cells in the spleen of mice treated with LPS or NaCl 2 weeks earlier. (**B**) Flow-cytometric analysis of B cells in spleens of Rag2^−/−^ mice treated with LPS or NaCl 2 weeks earlier, and reconstituted with GFP^+^ B cells before LPS/NaCl. (**C**) CD19^+^ B cells per femur of mice treated with NaCl or LPS 2 weeks earlier. (**D**) Flow-cytometric analysis of FO, MZ and B1/IRA B cells in spleens of mice treated with LPS or NaCl 2 weeks earlier. (**E**) Gating strategy for splenic B cell subsets. (**F**) Flow-cytometric analysis of Pre B, iB/tB and B1/IRA B cells in the bone marrow of mice treated with LPS or NaCl 2 weeks earlier. (**G**) IgM plasma levels in NaCl or LPS pretreated uninfected mice and 18h p.i. with *E. coli*. (**H**) ASAT plasma levels of NaCl or LPS pretreated WT and sIgM^−/−^ mice 18h p.i. with *E. coli*. Data in (A) and (C-F) and are representative out of 2-3 experiments (n= 3-8/experimental group). Data in (G-H) are pooled from 2 independent experiments (n = 3-7/experimental group). Data in (B) are from a single experiment (n = 7/group) and all data are and presented as mean +/− SEM. * p ≤ 0.05 and ** p ≤ 0.01.

B1 B and IRA B cell-derived IgM was shown earlier to exert tissue protective properties and has been proposed as a possible mechanism of disease tolerance (18–24). In line with this, we found elevated plasma IgM levels in tolerant mice prior to infection, which – in contrast to sensitive, control animals – returned to baseline during sepsis, indicating LPS-induced induction of IgM, and consumption of IgM during sepsis in tolerant animals (Figure 3G). We therefore tested if IgM was an essential soluble mediator responsible for the tissue protection in tolerant mice. Unexpectedly though, mice lacking soluble IgM developed less severe organ damage, and LPS-pretreatment still induced tissue tolerance during sepsis (Figure 3H, Figure S3D). Taken together, tissue tolerance was associated with long-term changes in the B cell compartments in the spleen and bone marrow, and the B cells’ tissue protective effects in tolerant mice occurred in an IgM-independent manner.

### B cells impact neutrophils, the key effectors driving sepsis-induced tissue damage

Having determined the importance of B cells in mediating tissue tolerance during sepsis, and having ruled out B-cell mediated inflammation or IgM effects as driving forces, we wondered if B cells might impact on the functionalities of other immune effector cells in sepsis. To assess the tissue damaging potential of candidate effector cells in our model, we depleted neutrophils (ΔPMN) (Figure S4A), platelets (ΔPlt) (Figure S4B) or monocytes and macrophages (ΔM) (Figure S4C) in sensitive and tolerant mice, respectively, prior to *E. coli* infection. Surprisingly, monocyte and macrophage depletion neither influenced sepsis-induced tissue damage in sensitive animals nor did it impact on LPS-induced tissue tolerance (Figure 4A). Platelet or neutrophil depletion, in contrast, already exerted tissue protective effects in both groups, illustrated by greatly reduced ASAT levels in sensitive and tolerant mice (Fig. 4A). However, while LPS-pretreatment still enhanced tissue tolerance in ΔPlt mice, it did not result in any additive beneficial effects in ΔPMN animals (Figure 4A), similar to what we had observed upon B cell deficiency (Figure 2D). These data support the reported role of platelets and neutrophils in promoting tissue damage during sepsis (4, 5) and proved neutrophils to be key effector cells of tissue protection in tolerant animals. Of note, no significant impact on the pathogen load was detectable in any of the groups (Figure S4D).

**Figure 4:**
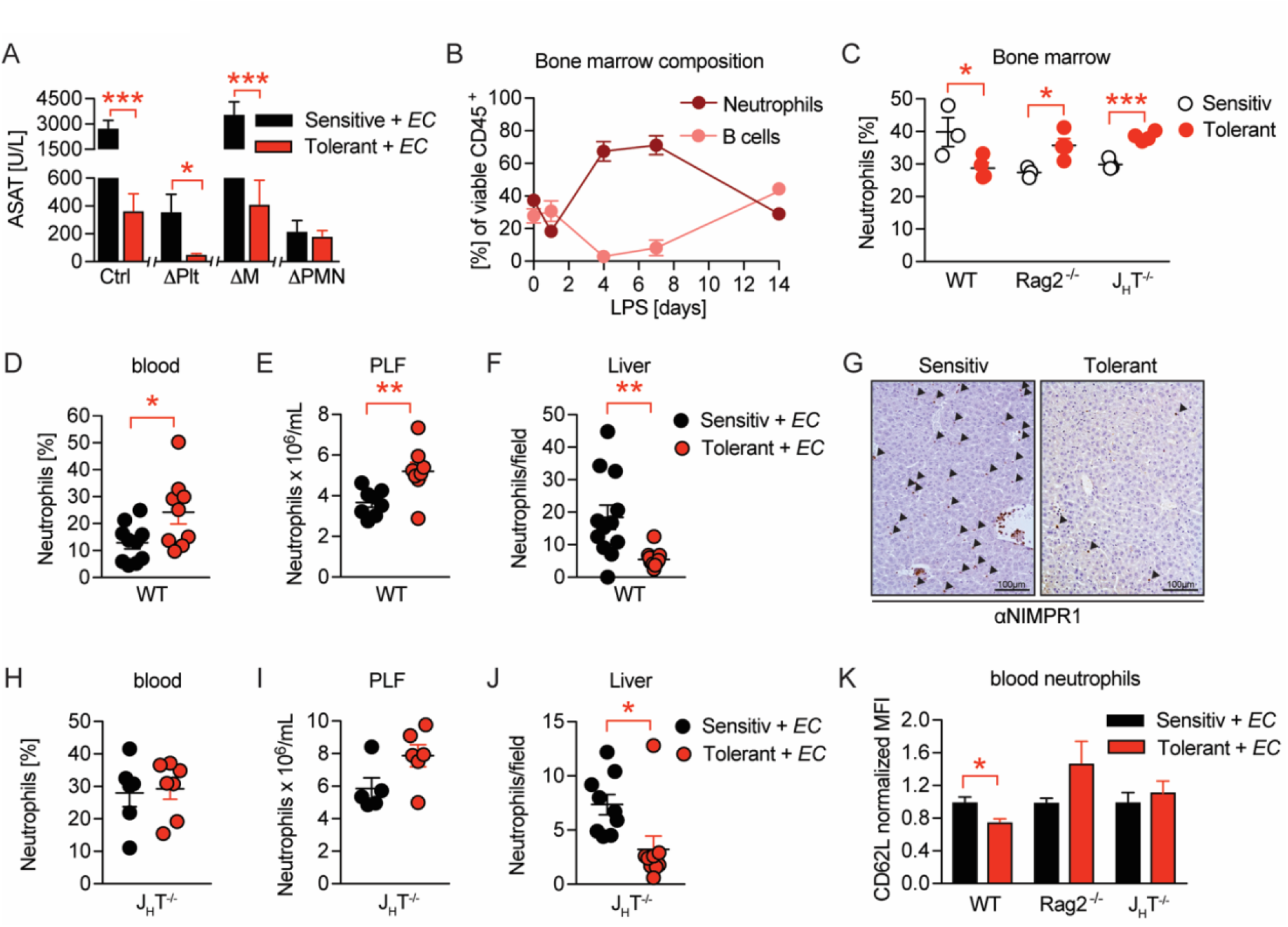
B cells impact neutrophils, the key effectors driving sepsis-induced tissue damage. (**A**) ASAT plasma levels 18h p.i. with *E. coli* in LPS or NaCl pretreated mice, in which platelets, monocytes/macrophages or neutrophils, respectively, were depleted before infection. (**B**) Flow-cytometric analysis of bone marrow neutrophils and B cells after i.v. administration of LPS at time = 0h. (**C**) Flow-cytometric analysis of neutrophils in the bone marrow of wildtype, Rag2^−/−^ and J_H_T^−/−^ mice 2 weeks after LPS or NaCl treatment. (**D**-**E**) Flow-cytometric analysis of neutrophils of wildtype mice pre-treated with NaCl or LPS, respectively, and infected for 18h with *E. coli*, in blood (D) and PLF (E). (**F**-**G**) Quantification of (F) immunohistological staining for NIMP-R1^+^ cells on liver sections (G) of mice pre-treated with NaCl or LPS, respectively, and infected with *E. coli* for 18h. (**H**-**J**) Flow-cytometric analysis of neutrophils 18h p.i. with *E.* coli in blood (H), PLF (I) and liver (J) of J_H_T^−/−^ mice. (**K**) Flow-cytometric analysis of blood neutrophil CD62L expression of WT, Rag2^−/−^ and J_H_T^−/−^ mice at 18h p.i. with *E. coli*. Data in (A) shown for the control group and neutrophil depletion are pooled from 2 independent experiments (n = 4-6/experimental group), platelet and monocyte/ macrophage depletion represent a single experiment (n = 8/group). Data in (B), (D-E), (F) and (J) are pooled from 2 independent experiments (n = 4-8/experimental group). Data in (H-I) are representative of 2 experiments (n = 5-8/group). Data in (K) are from a single experiment (n = 4-8/group). All data are presented as mean +/− SEM. * p ≤ 0.05, ** p ≤ 0.01 and *** p ≤ 0.001.

Considering our observation of LPS induced organ protection in a B cell and neutrophil dependent manner, we hypothesized an alliance between neutrophils and B cells in regulating tissue tolerance during sepsis. In steady state, up to 70% of CD45^+^ bone marrow cells are composed of B cells and neutrophils, where both populations constitutively reside and mature by sharing the same niche (25). We therefore first analyzed bone marrow B cell and neutrophil dynamics after LPS challenge, and discovered substantial stress-induced granulopoiesis, peaking around day four post LPS exposure, while B cells were regulated in a reciprocal fashion as they vanished by day four post LPS injection, to then increase and remain elevated two weeks post LPS treatment (Figure 4B and 3C) in tolerant, as compared to sensitive animals. At the same time total and relative neutrophil numbers in the bone marrow remained slightly reduced in tolerant mice, but elevated in the absence of B cells (Figure 4C and S4E).

Next, we assessed differences between sensitive and tolerant animals in infection-induced peripheral neutrophil migration and abundance, depending on the presence or absence of lymphocytes or B cells, respectively. While tissue tolerance was associated with elevated neutrophil abundance in blood and PLF of septic wild type mice (Figure 4D and 4E), neutrophil extravasation into tissues such as the liver and lung were substantially reduced (Figure 4F–G and S4F). In both Rag2^−/−^ and J_H_T^−/−^ mice, LPS pretreatment did not cause increased blood neutrophils nor a significant accumulation in the PLF during sepsis (Figure 4H–I and S4G–H). However, infection-induced neutrophil migration into liver tissue was still reduced after LPS pretreatment in J_H_T^−/−^ mice (Figure 4J), but not in Rag2^−/−^ animals (Figure S4I). This suggested that in tolerant animals, B cells might affect systemic neutrophil trafficking and turnover after LPS-pre-exposure, whereas the suppressed neutrophil extravasation to the livers of tolerant mice occurred independent of B cells. In support of this idea, we discovered that blood neutrophils of tolerant mice expressed lower CD62L levels upon infection than those of sensitive controls, and that this phenotype required the presence of B cells (Figure 4K). While CD62L has been studied extensively for its importance in neutrophil adhesion and rolling over the vascular endothelium (26), a recent study has identified decreased CD62L expression indicative of neutrophil aging, a process that is counteracted by Cxcr4 signaling, the master regulator of neutrophil trafficking between the bone marrow and the periphery (27–29). Based on these findings, we hypothesized that B cells might regulate sensitivity and tolerance during sepsis by affecting the functional plasticity and tissue damaging properties of neutrophils.

### Neutrophil tissue damaging properties are modulated by bone marrow B cells via Cxcr4

To assess the functional alterations in neutrophils, which confer tissue protection and tolerance during sepsis, we sorted neutrophils from the blood and bone marrow of sensitive and tolerant mice, i.e. 2 weeks post NaCl or LPS treatment but prior to infection, and performed RNA sequencing (Figure S5A). Despite the supposedly short life span of neutrophils, tolerant mice exhibited a sustained transcriptional reprogramming of the neutrophil pool. Principal component analysis of the 1000 most variable genes revealed clustering of neutrophils according to the site of sampling (Figure 5A), likely reflecting the heterogeneity of neutrophils in bone marrow versus mature cells in blood (30, 31). Samples further separated according to treatment (Figure 5A) and we identified a substantial number of tolerance associated, differentially expressed genes (DEGs) in both, bone marrow and blood neutrophils (Figure S5B-D, Supplementary Table 1 and 2). Gene ontology (GO) enrichment analysis of blood neutrophil DEGs exhibited an enrichment of genes associated with immunity and defense responses (Fig. 5B). Strikingly, bone marrow neutrophils of tolerant mice showed an enrichment of genes associated with cell migration, trafficking and chemotaxis (Figure 5C), such as genes involved in Cxcl12/Cxcr4 signaling including Cxcr4 itself (Figure S5E).

**Figure 5:**
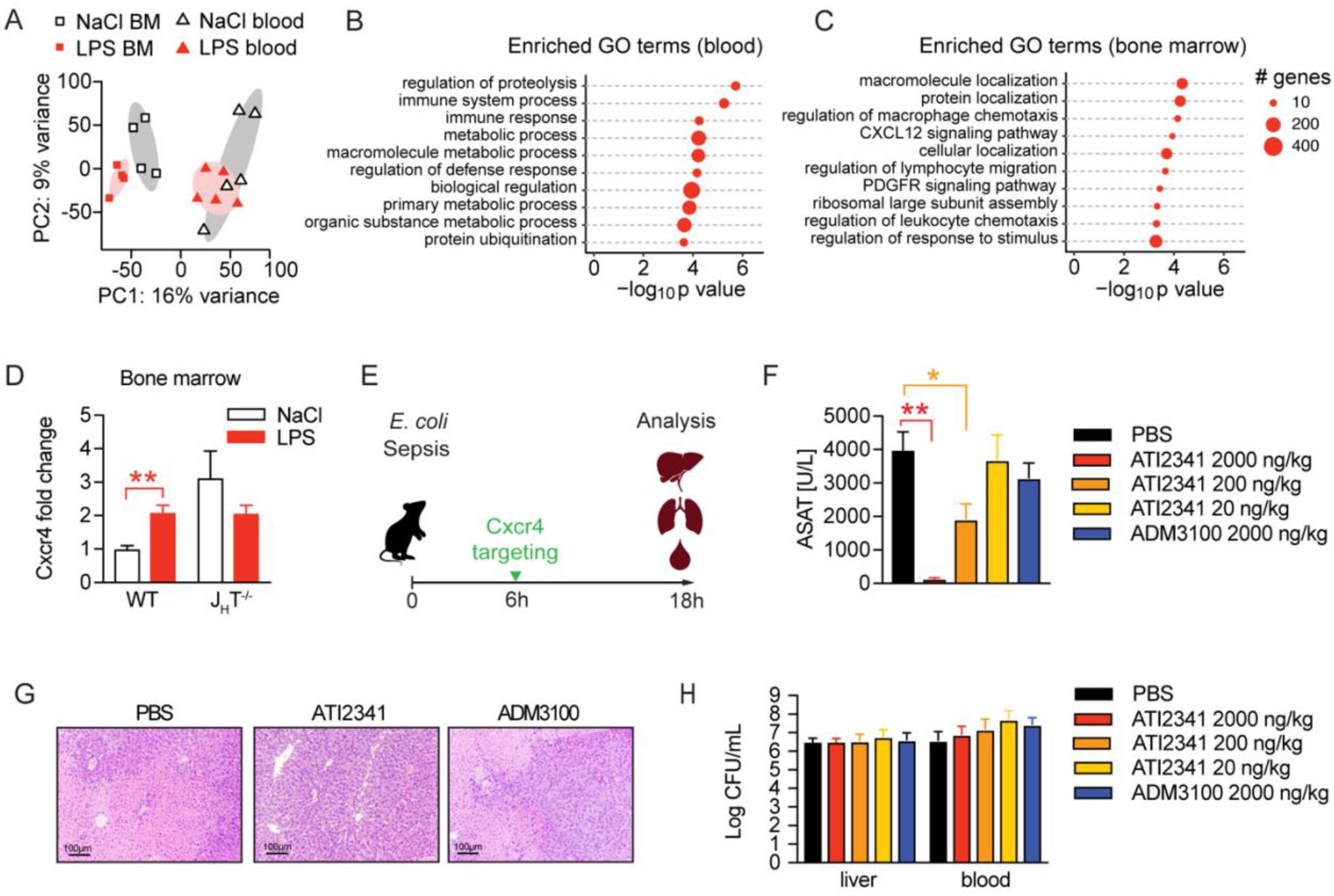
Neutrophil tissue damaging properties are modulated by bone marrow B cells via Cxcr4. (**A**) PCA of the top 1000 most variable genes expressed by neutrophils isolated from blood or bone marrow, of mice pretreated with LPS or NaCl 2 weeks earlier. (**B-C**) GO enrichment analysis of blood and bone marrow neutrophil DEGs. (**D**) *Cxcr4* mRNA expression in sorted bone marrow neutrophils from WT and J_H_T^−/−^ mice, pretreated with NaCl or LPS 2 weeks earlier. (**E**) Schematic depiction of the treatment procedure for the therapeutic application of a Cxcr4 agonist (ATI2341) and a Cxcr4 antagonist (ADM3100) at indicated doses. (**F**) ASAT plasma levels of mice 18h p.i. with *E. coli*, which were treated with depicted doses of Cxcr4 ligands (agonist ATI2341 or antagonist ADM3100, respectively) 6h p.i. (**G**) Representative liver histology (H&E stain) of mice 18h p.i. with *E. coli,* treated with PBS or the indicated Cxcr4 ligands (agonist ATI2341, antagonist ADM3100, at 2000ng/kg) 6h p.i. (**H**) Liver and blood CFUs of mice 18h p.i. with *E. coli* of mice, which were i.v. treated with depicted doses of Cxcr4 ligands (agonist ATI2341 and antagonist ADM3100, respectively) at 6h p.i. Data in (A-C) are from a single experiment (n = 4-5/group). Data in (D) are from an independent experiment (n = 3-4), versus data shown in (A-C). Data in (F-H) are representative of 2 independent experiments (n = 3-8/experimental group). All data are and presented as mean +/− SEM. * p ≤ 0.05 and ** p ≤ 0.01.

Considering the reported importance of Cxcr4 signaling in neutrophil trafficking between the bone marrow and periphery (27–29), we verified an upregulation of Cxcr4 on bone marrow derived neutrophils of tolerant mice compared to sensitive control mice on a transcriptional (Figure 5D) and protein level (Figure S5F). Importantly, this Cxcr4 induction depended on B cells, as Cxcr4 expression levels did not change in neutrophils isolated from LPS pre-exposed J_H_T^−/−^ mice (Figure 5D and S5F). Based on these findings and the recent observation that Cxcr4 deficient neutrophils promote aging and neutrophil-induced vascular damage (27), we hypothesized that B cells impact the life cycle of neutrophils by influencing neutrophil Cxcr4 signaling, which in turn might promote tissue tolerance during a subsequent sepsis. We therefore tested whether targeting Cxcr4 would be sufficient to induce tissue tolerance during sepsis and treated mice with increasing doses of the Cxcr4 pepducin agonist ATI2341, or a well-established dose of the Cxcr4 antagonist AMD3100 (Figure 5E). Strikingly, administration of the Cxcr4 agonist ATI2341 prevented sepsis-induced tissue damage in a dose dependent manner, whereas blocking Cxcr4 had no impact (Figure 5F). Liver histology reflected the tissue protective effects of ATI2341 treatment, while control and ADM3100 treated mice developed profound liver necrosis (Figure 5G). At the same time, none of these treatments altered the bacterial load (Figure 5H), suggesting that activation of Cxcr4 during sepsis induced tissue tolerance.

Taken together, by studying a model of tissue tolerance during sepsis, we here revealed a crosstalk between neutrophils and B cells in the bone marrow, in which B cells influence neutrophils likely by modulating Cxcr4 related pathways. In line with this idea, we found that administration of a Cxcr4 agonist limited tissue damage during severe sepsis, indicating that Cxcr4 signaling restrains the tissue damaging properties of neutrophils during infection.

## Discussion

Sepsis-induced tissue and organ damage are severe clinical complications responsible for the high fatality rate in patients suffering from sepsis (1, 3). To this end, no therapeutic option exists that can successfully prevent organ failure in septic patients. We established and investigated a model of disease tolerance during sepsis, which enabled us to reveal the importance of B cells and neutrophils in mediating tissue tolerance in the context of severe infections. Building on the reported interplay between these two cell populations, our data suggest that B cells shape neutrophils’ tissue damaging properties by modulation of neutrophil Cxcr4 signaling. Targeting Cxcr4 using a pepducin agonist protected mice from tissue damage during sepsis without affecting the bacterial load, indicating a Cxcr4-dependent disease tolerance mechanism.

Cellular depletion strategies allowed us to distinguish the contribution of selected cell types to infection-associated tissue damage in sepsis and, in parallel, to study their involvement in tissue tolerance mechanisms. The depletion of neutrophils as well as the absence of B cells fully abrogated tissue damage during primary sepsis (i.e. without prior tolerance-induction by LPS exposure), pointing towards a common, deleterious axis of these two immune cell types. It is well established that protective neutrophil effector functions during infection can be accompanied by severe collateral damage due to their tissue damaging properties by releasing inflammatory mediators such as IL-1β (32) and reactive oxygen species (33), or via tissue-factor mediated activation of coagulation (34–36) and the release of neutrophil extracellular traps (NETs) (37, 38). Our data indicate, that neutrophils are the primary effector cells that drive tissue damage, while B cells impact tissue damage by modulating neutrophil effector functions. It was demonstrated earlier that mature, splenic B2 cells promote neutrophil activation and their subsequent tissue damaging properties by boosting type-I IFN dependent early inflammation during polymicrobial sepsis (13). In support of proinflammatory, tissue-damaging properties of mature B2 cell subsets, we found splenectomy similarly protective as B cell deficiency during primary sepsis and reconstitution of Rag2^−/−^ mice with B cells to increase tissue damage. Interestingly, we did not identify an important role for the proposed IFNAR-driven inflammatory function of B cells (13) in sepsis, and inflammation did not differ between wild type and lymphocyte deficient mice. However, it seemed counterintuitive at first, that the absence of neutrophils or B cells, respectively, prevented tissue damage in a primary infection, while they at the same time seemed critical for tissue protection in a model of LPS-induced tolerance. We thus hypothesized that B1 and B1-like cells, in contrast to B2 cells, reduced neutrophil’s tissue damaging effector functions. Using sIgM deficient mice, enabled us to rule out a major role for IgM in tissue tolerance during sepsis, even though IgM was reported to exhibit anti-thrombotic functions in cardiovascular diseases (39) and high plasma IgM levels positively correlate with a better outcome in human sepsis (24) and mouse models (23). However, while sIgM deficiency did not prevent LPS-induced tolerance, naïve sIgM^−/−^ mice developed less organ damage during primary sepsis as compared to control animals. As sIgM deficiency goes along with a decreased abundance of B2 and an increased abundance of B1 cells (40) this further supported the notion of tissue damaging B2, and tissue protective B1 cells.

Since we discovered that LPS-induced protection was still observed in splenectomized animals, we considered the possibility that B cells regulate infection-induced neutrophil functionalities via effects exerted by sharing the same bone marrow niche. In fact, B cells, neutrophils and their precursors build up the majority of the constitutive CD45^+^ bone marrow cell pool, where they mature while sharing the same niche (25). Due to their potential tissue damaging properties, granulopoiesis and neutrophil trafficking is tightly controlled. Under steady state conditions in mice, only 2% of mature neutrophils circulate through the body, while the majority of cells is stored in the bone marrow from where they can be quickly released upon e.g. infection, to traffic to the periphery (30). Accumulating evidence highlights the substantial plasticity and functional heterogeneity of neutrophils, dependent on their localization (41), circadian rhythm (27) or maturation stage (31).

It has been proposed earlier that neutrophils and B cells regulate each other in a reciprocal fashion in the bone marrow (42). Based on our finding of a long-lasting transcriptional reprogramming in the neutrophil compartment and B cell dependency of tolerance-associated Cxcr4 upregulation, it is tempting to speculate that B cells act as important regulators of granulopoiesis and neutrophil trafficking at steady state and under inflammatory conditions. Cxcr4 interaction with its ligand Cxcl12 (stromal cell-derived factor 1, SDF1) has been shown to be critical for neutrophil release to the periphery as well as their homing back to the bone marrow when they become senescent (28, 29). Importantly, Cxcr4 signaling is essential, as *Cxcr4* knockout mice die perinatally due to severe developmental defects ranging from virtually absent myelopoiesis and impaired B lymphopoiesis to abnormal brain development (43). Antagonizing SDF1/Cxcr4 signaling is approved for stem cell mobilization from the bone marrow and is under extensive research in oncology, as it is critical for tumor development, metastasis and tumor cell migration (44). More recently, Cxcr4 signaling was described to delay neutrophil aging and to protect from vascular damage in an ischemia reperfusion model (27), supporting our data showing the tissue protective effects of upregulated Cxcr4 on neutrophils in sepsis. Strikingly, activating, but not antagonizing, Cxcr4 during sepsis induced tissue tolerance, suggesting that B cell driven regulation of Cxcr4 is a potential mechanism of disease tolerance and thus might be an interesting therapeutic target during severe sepsis.

## Material and Methods

### Reagents and recources

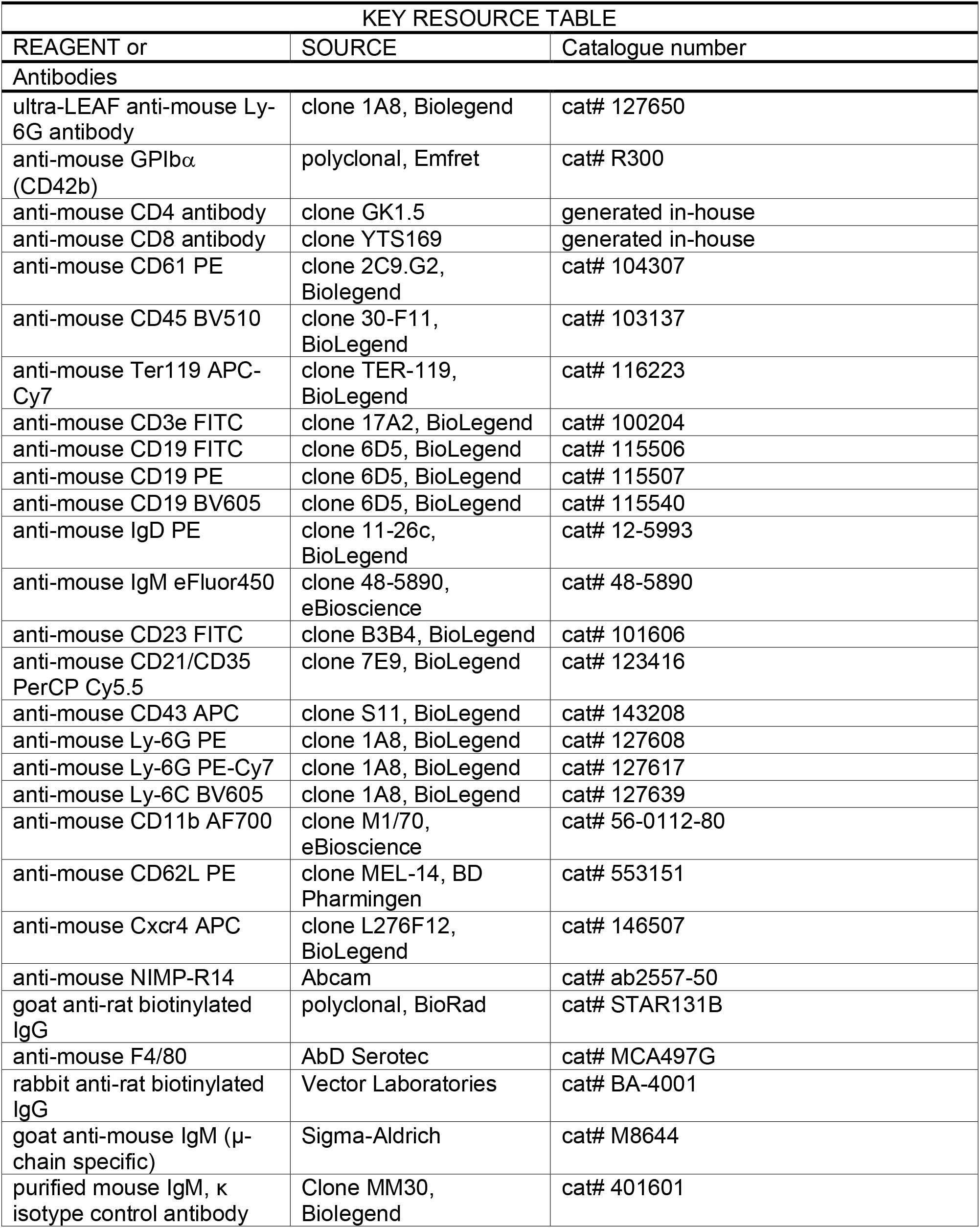
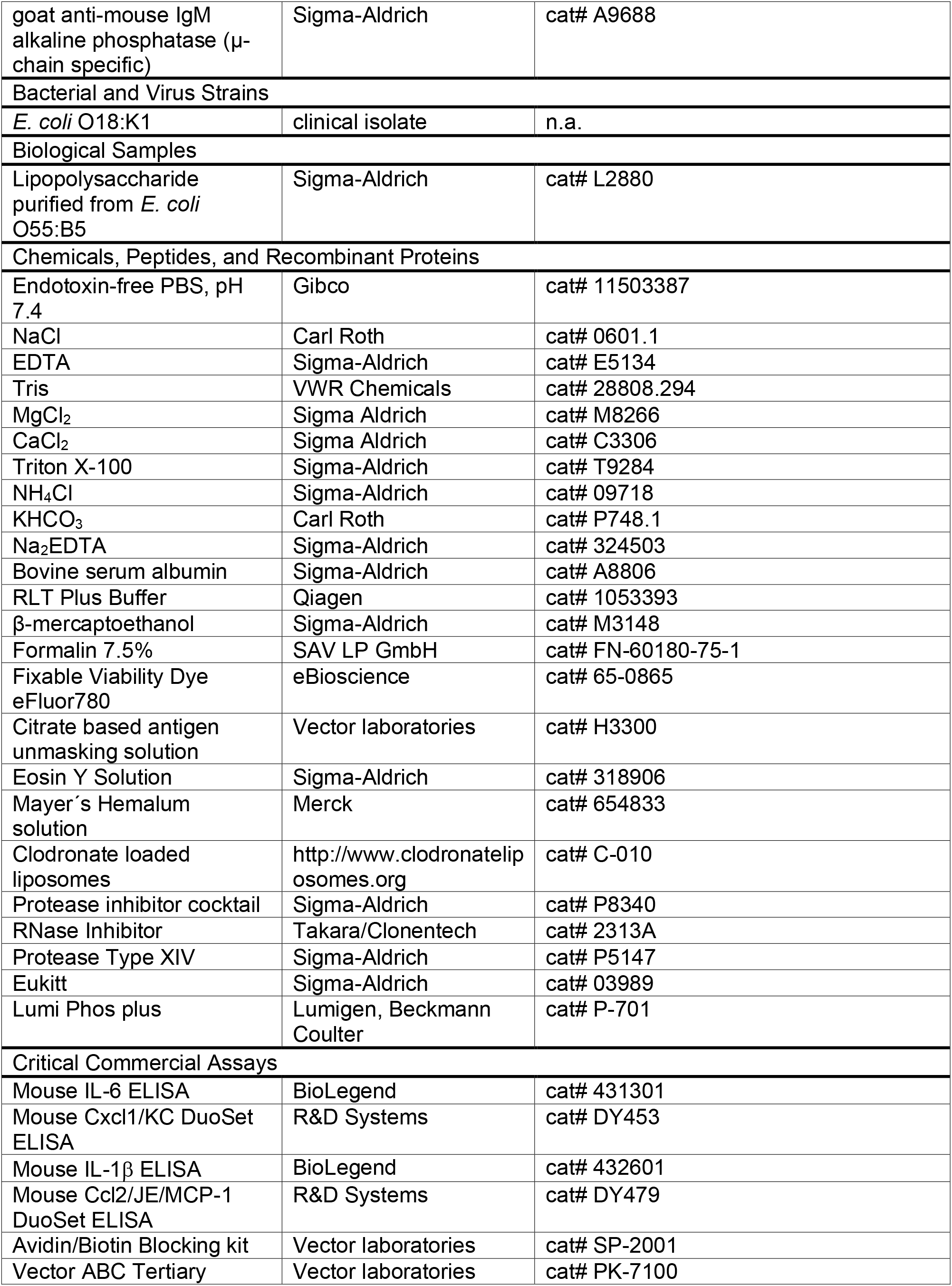
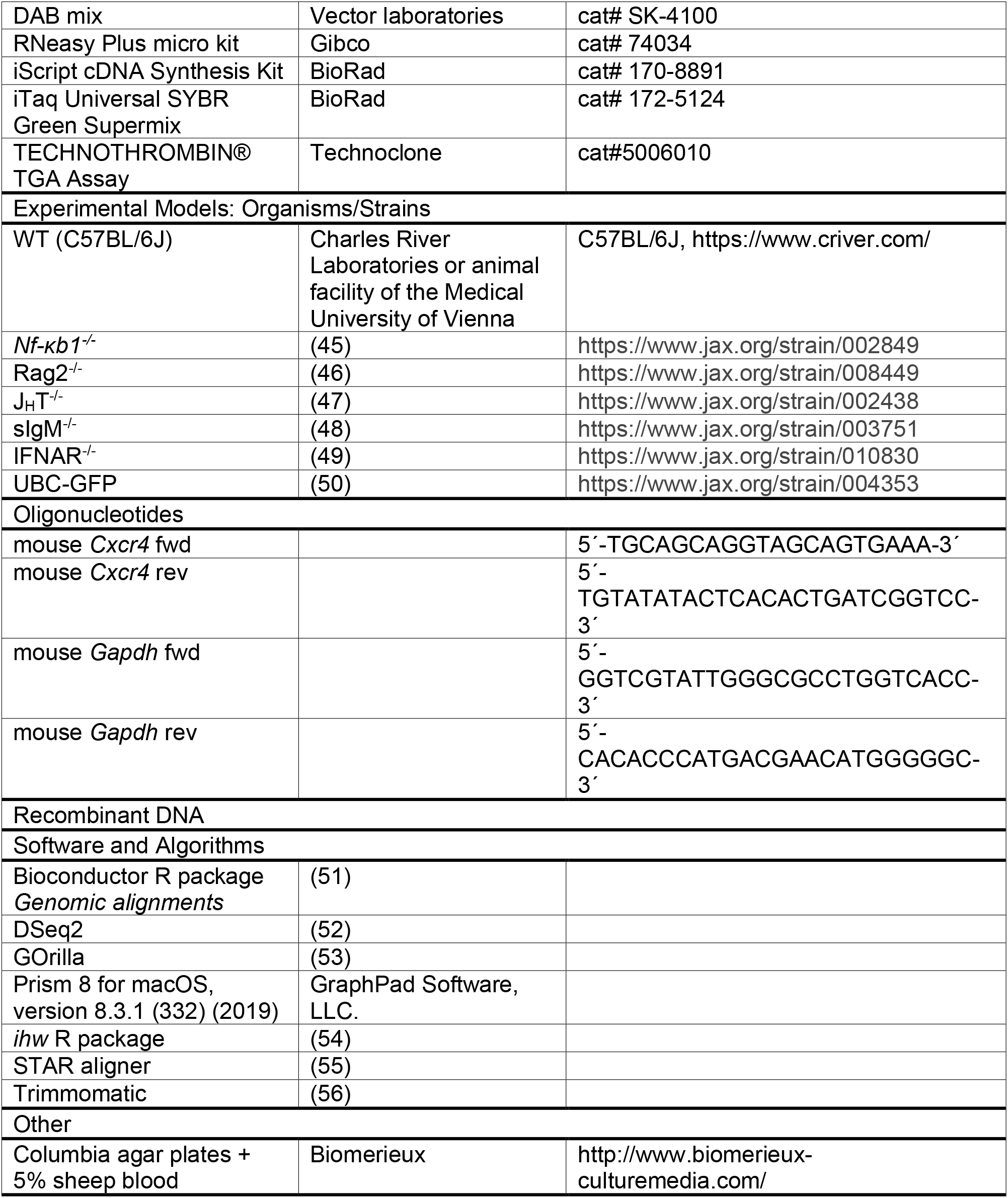

## Methods

### Animal studies

All experiments were performed using age-matched 8 to 12-week-old female mice. Wild type C57BL/6 mice were obtained from Charles River Laboratories or bred in the animal facility of the Medical University of Vienna. *Nf-κb1*^−/−^ mice were kindly provided by Derek Mann (Newcastle University, UK). Rag2^−/−^, J_H_T^−/−^, sIgM^−/−^, UBC-GFP, and IFNAR1^−/−^ mice were bred in the animal facility of the Medical University of Vienna.

All *in vivo* experiments were performed after approval by the institutional review board of the Austrian Ministry of Sciences and the Medical University of Vienna (BMWF-66.009/0272-II/3b/2013 and BMWF-66.009/0032-V/3b/2019).

### Mouse model of tolerance to E. coli peritonitis

Tolerance was induced by i.v. injection of 30μg *E. coli* LPS (Sigma-Aldrich) at indicated times before intraperitoneal (i.p.) infection with 1-2×10^4^ *E. coli* O18:K1. *E. coli* peritonitis was induced as described previously (57–59). Mice were humanely killed at indicated time points and blood, peritoneal lavage fluid (PLF) and organs were collected for further analysis. Peritoneal cell numbers were determined with a hemocytometer, and cytospin preparations were stained with Giemsa for differential cell counts and/or flow cytometry. Organs were stored in 7.5% formalin for histology or homogenized using Precellys 24 (Peqlab Biotechnologie GmbH) and prepared for further analysis as described earlier in detail (60). For ELISA, lysates were incubated in Greenberger lysis buffer (300mMol NaCl, 30mMol Tris, 2mMol MgCl_2_, 2mMol CaCl_2_, 1% Triton X-100, 2% protease Inhibitor cocktail) (61), and supernatants were stored at −20°C. For RNA isolation, lysates were stored in RLT buffer (Qiagen, containing 1% β-mercaptoethanol) at −80°C. Pathogen burden was evaluated in organ homogenates by plating serial dilutions on blood agar plates (Biomerieux), as previously described (57). Blood platelet counts were determined in freshly isolated anticoagulated EDTA blood using a VetABC differential blood cell counter. Liver transaminase levels (ASAT, ALAT) were quantified in the plasma using a Cobas c311 analyzer (Roche). IL-1, IL-6, TNF, CCL2 and CXCL1 were quantified using commercial ELISA kits according to manufacturers’ instructions. IgM levels were detected by coating plates with an anti-mouse IgM capture antibody (Sigma-Aldrich), followed by blocking with 1% BSA in PBS (containing 0,27mM EDTA) and incubation with plasma samples and standard dilutions of control mouse IgM (BioLegend). After several washing steps with PBS/EDTA, plates were incubated with an alkaline phosphatase-conjugated goat anti-mouse IgM (Sigma), washed with TBS (pH 7,4) and chemiluminescence was developed using Lumi Phos Plus (Lumigen) reagent.

### Cell depletions

Neutrophil or platelet depletion was achieved by i.v. injection of depletion antibodies 36h prior induction of *E. coli* peritonitis. Neutrophils were targeted using ultra-LEAF anti Ly-6G antibody (1mg/mouse, Biolegend) and platelets by injection of anti GPIbα (CD42b, 40μg/mouse) (Emfret). CD4^+^ and CD8^+^ T cell depletion was performed by i.v. administration of anti CD4 (200μg/mouse) or anti CD8 (400μg/mouse) antibodies 36h prior LPS treatment and repeated every three days until sepsis was induced by *E. coli* injection. Monocytes and macrophages were depleted by single i.v. administration of Clodronate loaded liposomes. Depletion of platelets, T- and B cells was verified by flow-cytometry. Neutrophil depletion was confirmed by differential cell counts of Giemsa stained PLF cytospins and macrophage depletion by immunohistochemistry for F4/80^+^ cells on formalin fixed liver sections.

### Cell transfers and splenectomy

Splenocytes were isolated from naïve WT C57BL/6 mice and i.v. injected into Rag2 deficient mice (1 × 10^7^ cells/mouse) after erythrocyte lysis using ACK lysis buffer (150mM NH_4_Cl, 10mM KHCO_3_, 0.1mM Na_2_EDTA, pH 7.2 – 7.4). Four days later, transplanted animals were pretreated with NaCl or LPS and two weeks later, challenged with *E. coli* as described above. Resting B cells were isolated from spleens of naïve UBC-GFP mice using magnetic beads (Milteny Biotec, Mouse B cell isolation kit) and i.v. injected into Rag2 deficient mice (5 × 10^6^ cells/mouse) after erythrocyte lysis (ACK lysis buffer) two weeks and four days prior to LPS/NaCl treatment. After pretreatment with NaCl or LPS transplanted animals were challenged with *E. coli* as described above. Mice were splenectomized or sham operated as described previously (62) and after 1 week recovery, treated with NaCl/LPS and challenged with *E. coli* as described above.

### In vitro thrombin-generation assay

Thrombin generation was assayed according to the manufacturer’s instruction (Technoclone). Briefly, citrated platelet poor plasma was thawed and shortly vortexed, diluted 1:2 with PBS and transferred onto a black NUNC Maxisorp plate. Fluorogenic thrombin generation substrate containing 15mM CaCl_2_ was added and the plate immediately read for 60min with Excitation/Emission at 360nm/460nm. Values were automatically calculated by the provided software.

### Flow cytometry

Splenocytes were isolated by passaging spleens through 70μm cell strainers and after erythrocyte lysis, single-cell suspensions were obtained. Bone marrow cells were obtained by flushing femurs, followed by filtering through 70μm cell strainers. Cells were counted using a CASY cell counter and after unspecific binding was blocked using mouse IgG (Invitrogen), cells were stained in PBS containing 2% FCS using antibodies (see table) against mouse CD45, CD3, CD19, CD23, IgM, CD21, CD43, CD11b and Ly-6G. This was followed by incubation with a Fixable Viability Dye eFluor 780 (eBioscience) according to the manufacturer’s instructions to determine cell viability. After several washing steps, cells were fixed (An der Grub Fix A reagent) and analyzed via flow cytometry using a BD LSRFortessa™ X-20 cell analyzer.

### Cell sorting, RNA sequencing and RT-PCR

For RNA sequencing, 200 neutrophils (defined as single/live/CD45^+^/CD3^−^/CD19^−^/Ly6G^+^/Ly-6C^int+^) were sorted from mouse bone marrow single cell suspensions (prepared as indicated above) into 4μL cell lysis buffer containing nuclease-free H_2_O with 0.2% Triton X-100 (Sigma-Aldrich) and 2 U/μl RNase Inhibitor (Takara/Clonentech) using a FACSAria Fusion cytometer. Cell lysates were stored at −80°C. Library preparation was performed according to the Smart-Seq2 protocol (63), followed by sequencing of pooled libraries on the Illumina HiSeq 2000/2500 (50 bp single‐read setup) at the Biomedical Sequencing Facility of the Medical University of Vienna and CeMM. For analysis, reads were adapter-trimmed (Trimmomatic) (56) and aligned to the *mm10* reference genome (*STAR aligner*) (55). Counting of reads mapping to genes was performed using the *summarizeOverlaps* function (*Bioconductor* R package *GenomicAlignments*) (51). Differentially expressed genes were identified using *DESeq2* (52), whereby separate models per organ and condition (bone marrow or blood, respectively, +/− LPS or NaCl, respectively treatment) were formulated for all pairwise comparisons. Filtering was performed by independent hypothesis weighting (*ihw* R package) (54). Gene ontology (GO) enrichment analysis of neutrophil DEGs was performed using the GOrilla (Gene ontology enrichment analysis and visualization tool).

For verification of *Cxcr4* upregulation, 1.5×10^5^ neutrophils were sorted as described above into cold PBS containing 2% BSA, followed by centrifugation and resuspension of the pellet in 350μl RLT Buffer containing 1% (v/v) ß-mercaptoethanol. RNA isolation was performed using RNeasy Plus micro kit (Qiagen) according to the manufacturer’s instructions. Reverse transcription was performed using 150ng of isolated RNA and the iScript cDNA Synthesis Kit (Biorad), according to manufacturer’s instructions. Real-time PCR for mouse *Cxcr4* was performed with iTaq Universal SYBR Green Supermix reagents (Applied Biosystems) on a StepOnePlus Real-Time PCR System (Applied Biosystems) using *Gapdh* as a housekeeper.

### Histology

Liver sections (4 μm) were stained with H&E and analyzed by a trained pathologist in a blinded fashion according to a scoring scheme, involving necrosis, sinusoidal- and lobular inflammation, steatosis and endothelial inflammation (0 representing absent, 1 mild, 2 moderate, and 3 severe). The sum of all parameters indicated the total histology score. After staining for fresh fibrin (MSB stain, performed at the routine laboratory at Newcastle University), samples were scored for the presence of microthrombi by a trained pathologist in a blinded fashion. NIMPR1 immunostaining was performed on paraffin-embedded liver sections as described earlier (64). Briefly, antigen retrieval was achieved using a citrate-based buffer at pH 6.0 (Vector laboratories), followed by several blocking steps. Incubation with anti-NIMP-R14 antibody (Abcam) was performed at 4°C, over-night followed by 2h exposure to a biotinylated secondary goat anti-rat antibody (Serotec/Biorad). For F4/80 staining, antigen retrieval was achieved by protease type XIV (Sigma) digestion at 37°C for 20min, followed by several washings and blocking steps. After 1h incubation with an anti-F4/80 antibody (Serotec), exposure to a biotinylated secondary rabbit anti-rat antibody was performed at room temperature. Finally, both stains were incubated with VECTASTAIN Elite ABC Reagent and visualized using diaminobenzidine peroxidase substrate (Vector Laboratories), followed by counterstaining with hematoxylin and embedding (Eukitt, Sigma).

### Statistical analysis

Statistical evaluation was performed using GraphPad Prism software except for statistical analysis of RNA sequencing data, which was performed using R. Data are represented as mean ± SEM and were analyzed using either Student’s t-test, comparing two groups, or one-way ANOVA analysis, followed by Tukey multiple comparison test, for more than two groups. Differences with a p-value ≤ 0.05 were considered significant. For DEG, genes with an FDR-adjusted p value of < 0.1 were considered differentially expressed.

**Table 1.** DEG of blood neutrophils isolated from mice 2 weeks post NaCl or LPS treatment

**Table 2.** DEG of bone marrow neutrophils isolated from mice 2 weeks post NaCl or LPS treatment

## Author contributions

RG performed experiments, analyzed data and wrote the manuscript. BM and GO performed experiments and analyzed data. MLW, A-DG, AF, FQ, KL, AH, AK and AA helped with experimental procedures. IM, FOb and JGB performed pathological scorings. LB provided crucial reagents. FOa, IL and CJB contributed to the design of the study. SK conceived and supervised the project and wrote the manuscript.

## Acknowledgements

The authors thank Henriette Luise Horstmeier and Aysu Eshref for help with experimental procedures, Sophie Zahalka for help with illustrations and the animal caretakers at the Center for Biomedical Research, Medical University Vienna, for their expert work. We acknowledge the core facilities of the Medical University of Vienna, including the Flow Cytometry Facility and the Biomedical Sequencing Facility (BSF, jointly run with CeMM). This work was funded by the Austrian Science Fund (FWF) within the Doctoral Program Cell Communication in Health and Disease (CCHD, 1205FW), and the Special Research Projects Immunothrombosis (SFB 054-10) and Chromatin Landscapes (SFB 061-04) to S.K. FO is supported by MRC program Grants; MR/K0019494/1 and MR/R023026/1.

## Supplemental Figures

**Supplementary Figure 1:**
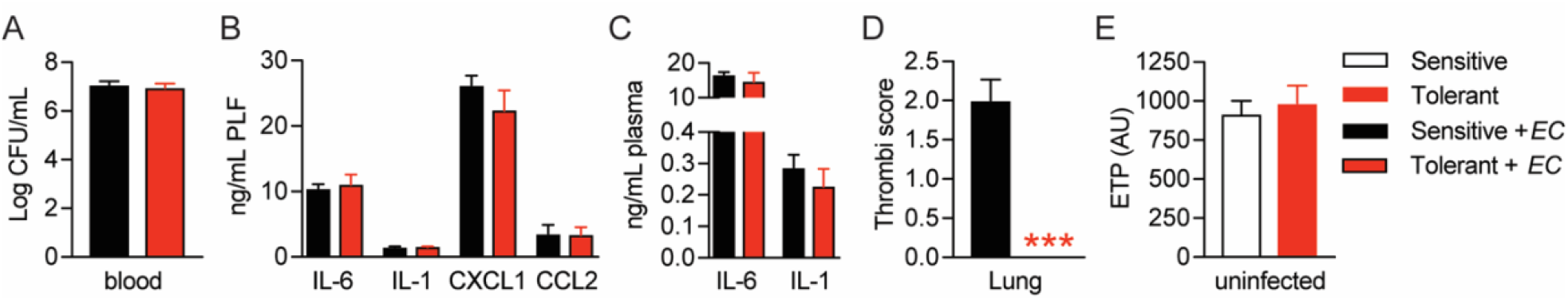
(**A**) Blood *E. coli* CFUs 18h p.i. with *E. coli* of mice, pretreated with LPS or NaCl 2 weeks before infection. (**B-C**) PLF and plasma cytokine and chemokine levels 18h. p.i. with *E. coli.* (**D**) Histopathological scoring of lung microthrombi 18h p.i. with *E. coli.* (**E**) Endogenous thrombin potential (ETP) of plasma samples from uninfected mice 2 weeks after LPS or NaCl pretreatment. Data depicted in (D) represent a single experiment (n = 8/group). All other data are representative for two or more independent experiments (n = 5-8/experimental group) and are and presented as mean +/− SEM. *** p ≤ 0.001.

**Supplementary Figure 2:**
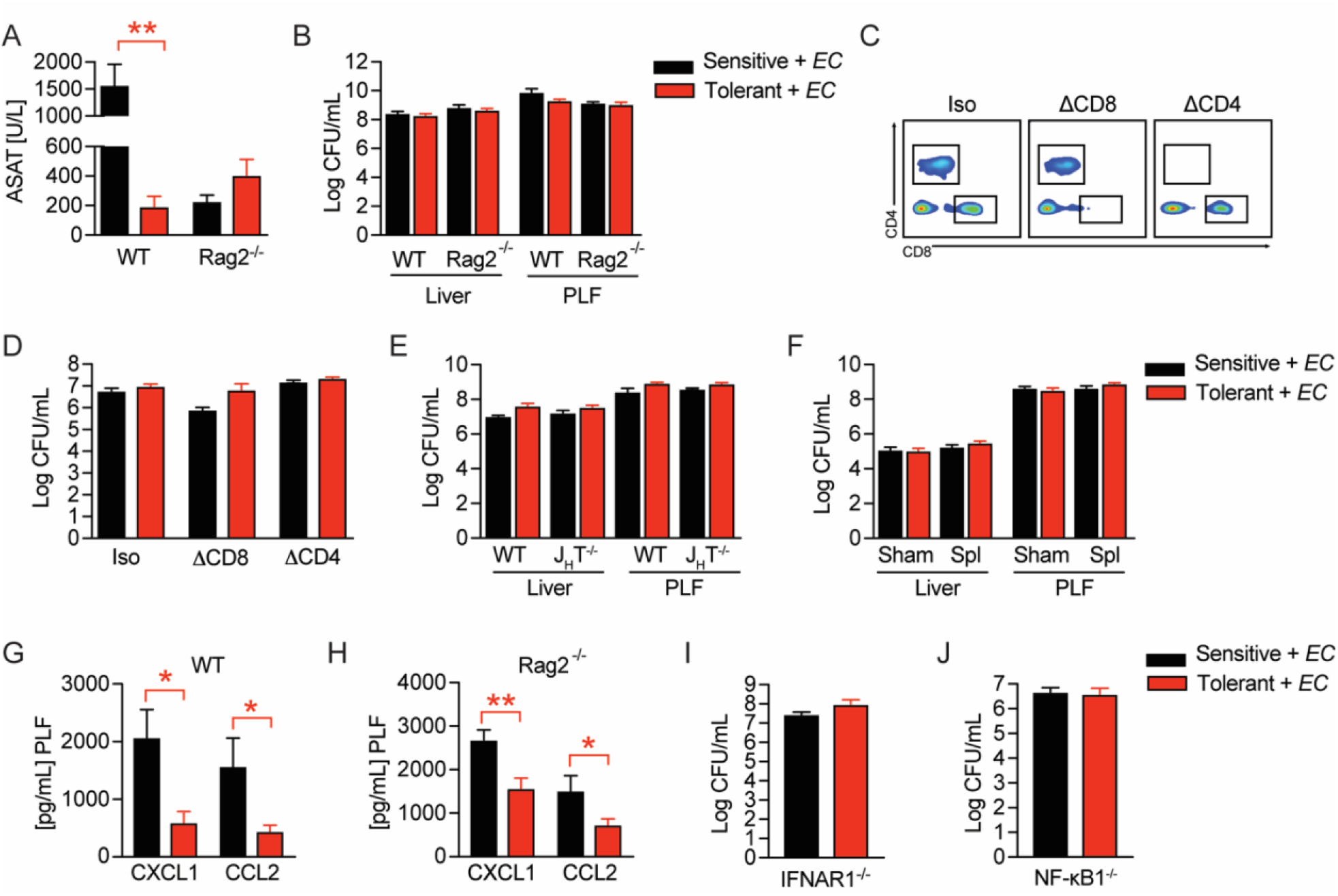
(**A**) ASAT plasma levels 18h p.i. with *E. coli* of NaCl or LPS pretreated WT and Rag2^−/−^ mice. (**B**) *E. coli* CFUs in livers and PLF of NaCl or LPS pretreated WT and Rag2^−/−^ mice 18h p.i. with *E. coli*. (**C**) Flow-cytometric analysis of CD4^+^ and CD8^+^ T cells in blood of mice after administration of anti-CD8, or anti-CD4, respectively, cell depletion antibodies. (**D**) *E. coli* CFUs in livers 18h p.i. with *E. coli* of mice, which received CD4^+^ or CD8^+^ T cell depleting antibodies before LPS or NaCl pretreatment. (**E**) *E. coli* CFUs in livers and PLF of NaCl or LPS pretreated WT and J_H_T^−/−^ mice at 18h p.i. with *E. coli*. (**F**) *E. coli* CFUs in livers and PLF 18h p.i. of mice, which were splenectomized or sham operated 1 week before LPS or NaCl pre-exposure (i.e. 3 weeks before bacterial infection). (**G**-**H**) CXCL1 and CCL2 levels in PLF of NaCl or LPS pretreated wildtype (I) or Rag2^−/−^ (J) mice 6h p.i. with *E. coli*. (**I**) *E. coli* CFUs in livers of NaCl or LPS pretreated IFNAR1^−/−^ mice 18h p.i. with *E. coli*. (**J**) *E. coli* CFUs in livers of NaCl or LPS pretreated NF-κB1^−/−^ mice 18h p.i. with *E. coli*. Data in (A-B) and (E-I) are representative out of 2-3 experiments (n = 4-8/experimental group). Data depicted in (C-D) are from a singl experiment (n = 6-8/group). Data in (J) are pooled from 2 experiments (n = 2-6/experimental group) and all data are presented as mean +/− SEM. * p ≤ 0.05 and ** p ≤ 0.01.

**Supplementary Figure 3:**
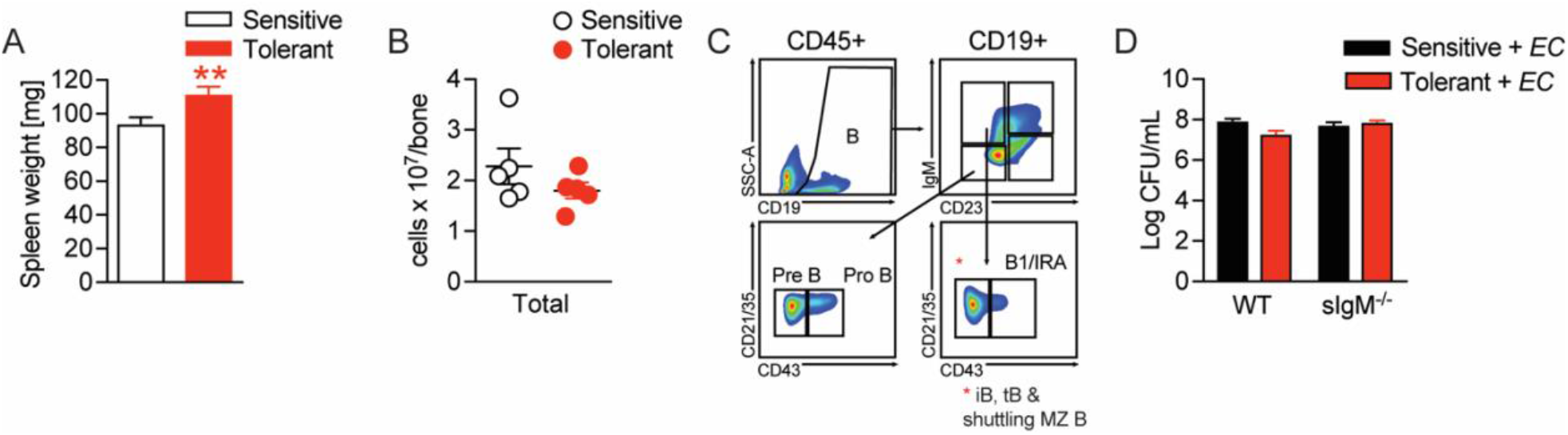
(**A**) Spleen weight of mice, which were treated with NaCl or LPS 2 weeks earlier. (**B**) Nucleated cells per femur of mice treated with NaCl or LPS 2 weeks earlier. (**C**) Gating strategy for bone marrow B cell subsets. (**D**) *E. coli* CFUs in livers of NaCl or LPS pretreated WT and sIgM^−/−^ mice 18h p.i. with *E. coli*. Data in (B) and (D) are representative out of 2-3 experiments (n = 4-8/experimental group). Data in (A) are pooled from 3 experiments (n = 2-6/experimental group) and all data are presented as mean +/− SEM. ** p ≤ 0.01.

**Supplementary Figure 4:**
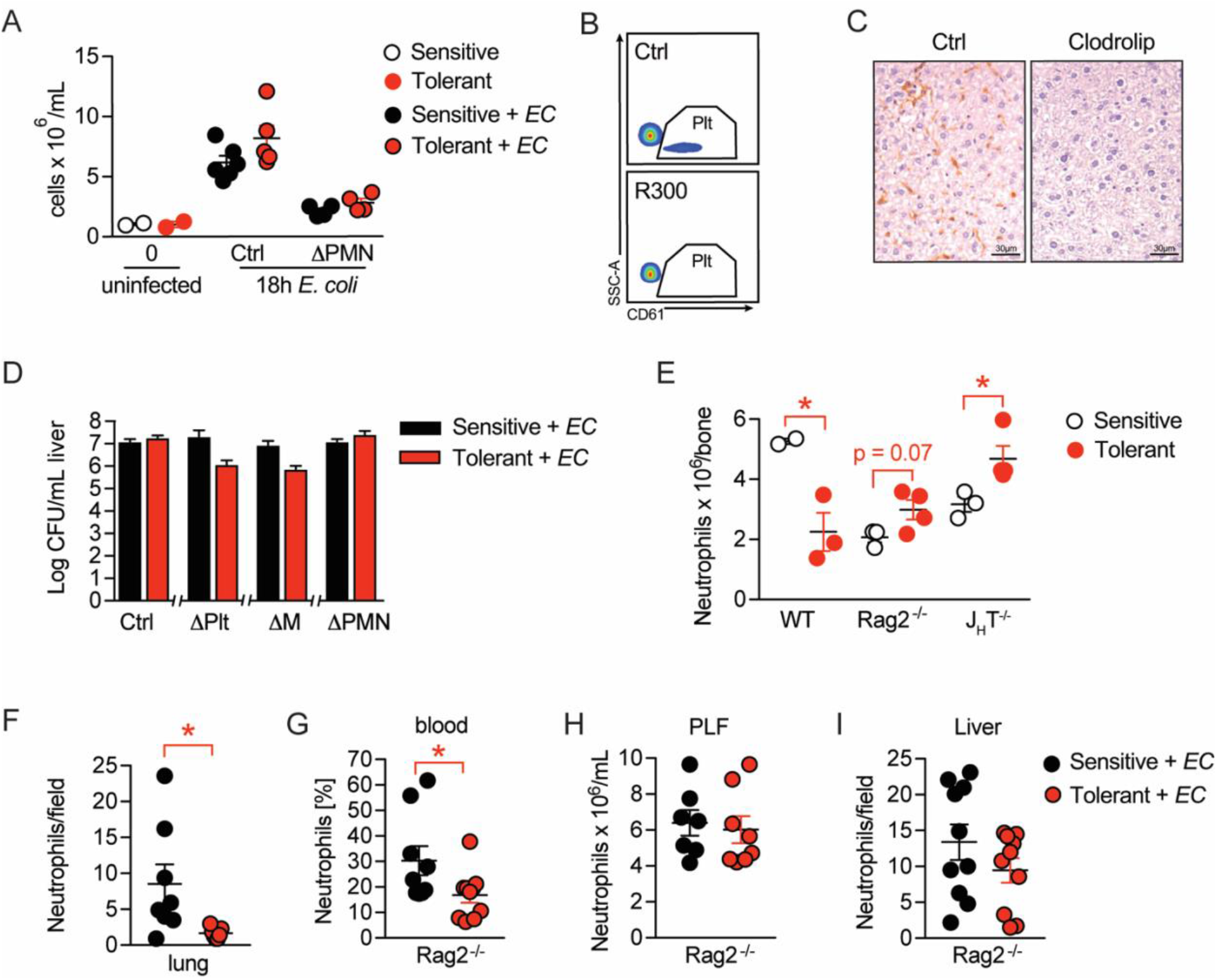
(**A**) PLF cell numbers of uninfected mice and mice 18h p.i. with *E. coli,* which received an anti-Ly-6G depletion antibody i.v. 24h before infection. (**B**) Flow-cytometric analysis of blood platelets (SSCA_low_, CD61^+^) in mice that received an anti-GPIbα (R300) depletion antibody 24h earlier. (**C**) Immunohistological staining for F4/80^+^ cells on liver sections of mice which received either clodronate-loaded liposomes (clodrolip) or empty control liposomes i.v. 24 hours earlier. (**D**) *E. coli* CFUs 18h p.i. with *E. coli* in livers of NaCl or LPS pretreated mice, which received depletion antibodies for platelets or neutrophils, or clodronate liposomes before infection. (**E**) Flow cytometric analysis of bone marrow neutrophil abundance 2 weeks after treatment with LPS or NaCl. (**F**) Quantification of NIMP-R1^+^ cells in immunohistological staining of lung sections from NaCl or LPS pretreated mice 18h p.i. with *E. coli*. (**G**-**H**) Flow-cytometric analysis of neutrophils in blood and PLF of NaCl or LPS pretreated Rag2^−/−^ mice 18h p.i.. with *E. coli*. (**I**) Quantification of NIMP-R1^+^ cells after immunohistological staining of liver sections from NaCl or LPS pretreated Rag2^−/−^ mice 18h p.i. with *E. coli*. Data shown in (A), (E) and (H) are representative out of 2 independent experiments (n = 2 -6 per experimental group for A and E and n = 7-8 per experimental group for H). Data in (B-C) are from a single experiment, which was set up to test the depletion efficiency in uninfected animals (n = 3 per experimental group). In (D) CFU data shown for the control group and neutrophil depletion are pooled from 2 independent experiments (n = 4-6 per experimental group) and data showing the platelet and monocyte/macrophage depletion are from a single experiment (n = 8 per group). Data depicted in (G) and (I) are pooled from 2 independent experiments (n = 4-6 per experimental group) and data in (F) are from a single experiment (n = 8 per group). All data are presented as mean +/− SEM. * p ≤ 0.05.

**Supplementary Figure 5:**
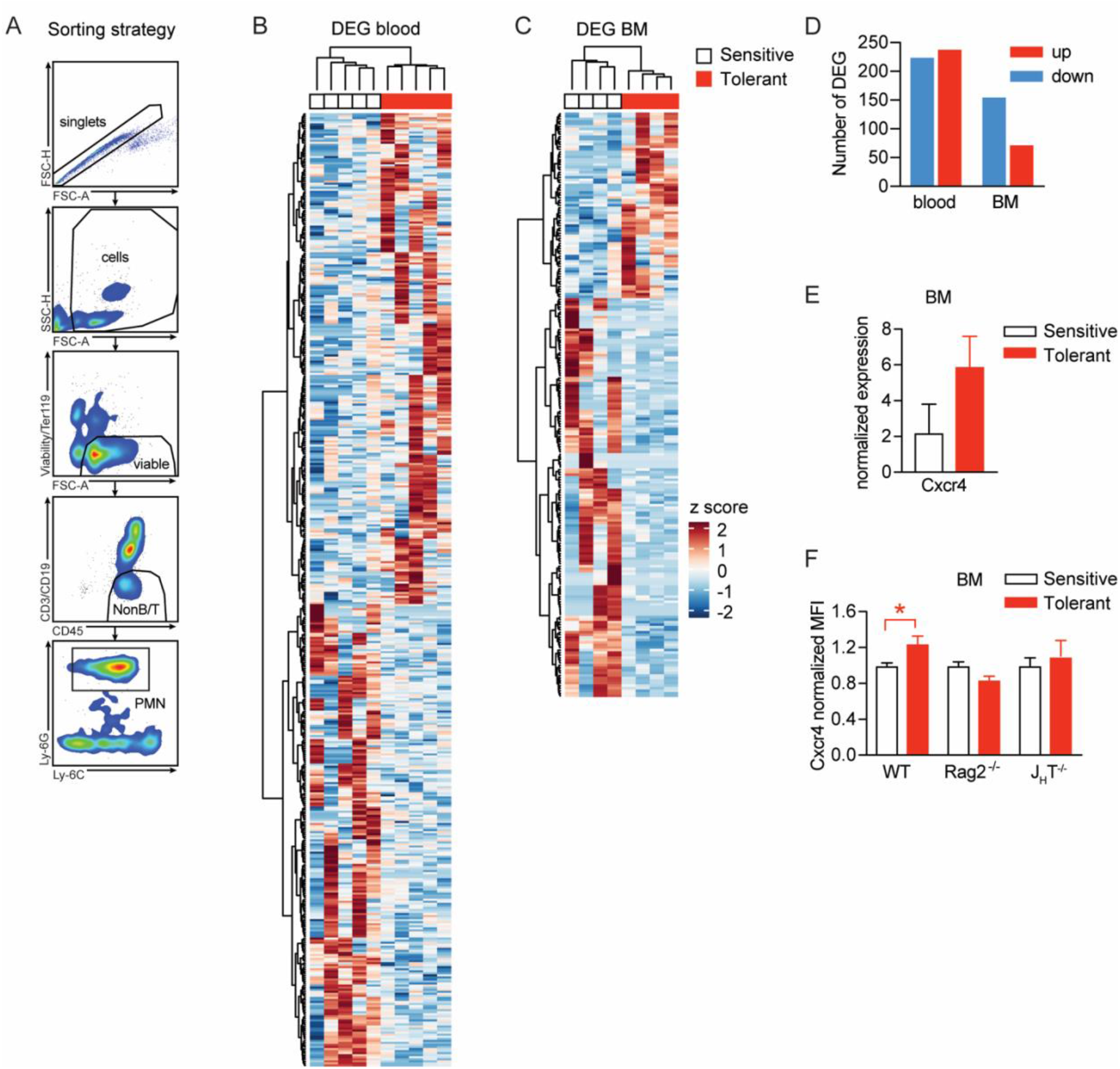
(**A**) Gating strategy applied for neutrophil sorting from blood and bone marrow. (**B-C**) Hierarchical clustering of DEG of blood and bone marrow neutrophils from mice that were pretreated with LPS versus NaCl 2 weeks earlier. (**D**) Number of up- and downregulated DEG in blood and bone marrow neutrophils from mice pretreated with LPS 2 weeks earlier, as compared to NaCl treated controls. (**E**) Normalized gene expression of *Cxcr4* in sorted bone marrow neutrophils. (**F**) Normalized MFI of Cxcr4 on bone marrow neutrophils assessed by flow cytometry in indicated mouse strains. Data in (A-E) are from a single experiment (n = 4-5/group) and data in (F) are from a different single experiment (n = 3-4/group). All data are presented as mean +/− SEM. * p ≤ 0.05.

## Abbreviations

ALAT: Alanine aminotransferase
ASAT: Apartate aminotransferase
Cxcr4: CXC-motive chemokine receptor 4
DEG: differentially expressed gene
*E. coli*: *Escherichia coli*
FO B cell: follicular B cell
IFNAR: interferon-α/β receptor
*i.p.*: intraperitoneally
IRA B cell: innate response activator B cell
*i.v.*: intravenously
LPS: Lipopolysaccharide
MZ B cell: marginal zone B cell
PLF: peritoneal lavage fluid
p.i.: post infection
TLR: Toll-like receptor

